# Shared neuronal bases of inhibition and economic choice in orbitofrontal cortex

**DOI:** 10.1101/2020.04.23.057455

**Authors:** Pragathi Priyadharsini Balasubramani, Benjamin Y. Hayden

**Author notes:** Corresponding Author: Pragathi Priyadharsini Balasubramani, Department of Psychiatry, University of California San Diego, La Jolla, CA. 92037. The authors declare no conflict of interest.

## Abstract

Economic choice and inhibition are two important elements of our cognitive repertoires that may be closely related. We and others have noted that during economic choice, options are typically considered serially; this fact provides important constraints on our understanding of choice. Notably, asynchronous contemplation means that each individual option is subject to an accept-reject decision. We have proposed that these component accept-reject decisions may have some kinship with stopping decisions. One prediction of this idea is that stopping and choice may reflect similar neural processes occurring in overlapping brain circuits. To test the idea, we recorded neuronal activity in orbitofrontal cortex (OFC) Area 13 while macaques performed a stop signal task interleaved with a structurally matched choice task. Using neural network decoders, we find that OFC ensembles have overlapping codes for stopping and choice: the decoder that was only trained to identify accept vs. reject trials performed with higher efficiency even when tested on the stop trials. These results provide tentative support for the idea that mechanisms underlying inhibitory control and choice selection may be subject to theoretical unification.

## INTRODUCTION

Foraging theory is a branch of behavioral ecology centered on understanding the psychology of reward-based choices in natural contexts (Stephens and Krebs 1986). It provides a unique perspective on many aspects of economic choice and has recently come to influence the neuroscience and psychology of decision-making (Pearson et al. 2014; Calhoun and Hayden 2015; Mobbs et al. 2018; Hayden et al., 2011). A core idea in foraging theory is that natural decisions are fundamentally structured around accepting vs. rejecting – taking or passing up a single option that is the sole or primary focus of attention (Shapiro et al. 2008; Vasconcelos et al. 2010; Kacelnik et al. 2011; Pirrone et al. 2017; Ojeda et al. 2018). Even ostensibly binary economic choices, in this view, reflect a pair of (potentially interacting) accept-reject choices. *Accepting* involves selecting the attended or activated option, or, more abstractly, performing the afforded action (Cisek and Kalaska 2010; Cisek and Pastor-Bernier 2014; Hayden and Moreno-Bote 2018). *Rejecting* involves countermanding the afforded action. An ostensibly binary economic choice, then, may be seen as two related decisions about whether to go or stop choosing the attended option or the afforded action (Krajbich et al. 2010; Kacelnik et al. 2011; Hayden 2018).

Here we consider one particular implication of this foraging-inspired line of work: that economic choice may be subject to theoretical unification with stopping decisions (Hayden 2018; Balasubramani et al. 2019). The rationale for this idea is straightforward. Both economic and stopping decisions involve a single option as an offer and a choice of whether to pursue that option or refrain from pursuing it. Both are framed around a default option (accepting and going, respectively) and a non-default (rejecting and stopping), that is, deliberate inhibition of the default option.

There is at least some evidence in support of this hypothesis. First, stopping and economic choice tend to activate a similar set of brain regions, including the pre-motor cortex, the ventrolateral prefrontal cortex, basal ganglia, and the thalamus (Schall et al. 2002; Aron and Poldrack 2006; Aron 2007; Sakagami and Pan 2007; Cisek and Kalaska 2010; Cisek 2012; Hampshire and Sharp 2015). Second, this way of looking at choice is consistent with some recent studies that suggest that binary choice involves a serial, not parallel, consideration of options (Krajbich et al. 2010; Strait et al. 2014; Rich and Wallis 2016; reviewed in Hayden and Moreno-Bote, 2018). These studies and others indicate that attention is largely limited to a single option, which is evaluated, often relative to the other one (Lim et al. 2011; Rudebeck and Murray 2014; Strait et al. 2014; Strait et al. 2015; Rich et al. 2017; Xie et al. 2018).

We recorded neuronal activity in orbitofrontal cortex (OFC) area 13. The importance of OFC for economic choice is largely undisputed, although its specific role remains to be determined (Wallis 2007; Schoenbaum et al. 2009; Padoa-Schioppa 2011; Rudebeck and Murray 2014; Wilson et al. 2014; Rich et al. 2017). It is clear, nonetheless, that activity of OFC correlates with the values of offers and of chosen options, and is likely to be critical for value comparison as well (Padoa-Schioppa and Assad 2006; Padoa-Schioppa 2013; Raghuraman and Padoa-Schioppa 2014; Wallis, 2007). In contrast to its clear role in choice, the contribution of the OFC to stopping remains to be determined. Specifically, some studies and surveys argue against its inhibitory role (Schoenbaum et al. 2003; Chudasama et al. 2006; Ghods-Sharifi et al. 2008; Rudebeck and Murray 2014; Stalnaker et al. 2015). Others argue in support of some inhibitory role (Mishkin 1964; Iversen and Mishkin 1970; Dias et al. 1996; Roberts and Wallis 2000; Horn et al. 2003; Eagle et al. 2007; Chikazoe et al. 2009; Majid et al. 2013; Bryden and Roesch 2015; Balasubramani et al. 2019). Our previous study on the topic suggests that it does contribute to inhibition/stopping, although this is part of its more complex role in setting the stage for action, which includes its economic functions (Balasubramani et al., 2019).

Here we sought to test one specific prediction of the overlap hypothesis by comparing neuronal activity in a choice task with that observed in a stopping task. For stopping task, we use the stop signal task framework, a standard tool for studies of stopping and inhibitory control (Logan and Cowan 1984; Logan 1994; Hanes and Schall 1995). We then designed a novel economic choice task that follows, as closely as possible the design of the stopping task and call it choice signal task, so that we could focus on differences relevant to our hypothesis.

## METHODS

Some of the data presented here were analyzed for a previous manuscript (Balasubramani et al., 2019). Specifically, data from the stopping task were analyzed in that manuscript. Data from the choice task were collected in the same sessions but not analyzed therein.

### Subjects

Two male rhesus macaques (Macaca mulatta, subject J, age 10, and subject T, age 5) served as subjects. All animal procedures were approved by the University Committee on Animal Resources at the University of Rochester and were designed and conducted in compliance with the Public Health Service’s Guide for the Care and Use of Animals.

### Recording site

A Cilux recording chamber (Crist Instruments) was placed over the Area 13 of OFC, as defined by (Paxinos and Watson 2006). The targeted area expands along the coronal planes situated between 28.65 and 33.60 mm rostral to the interaural plane with varying depth. Position was verified by magnetic resonance imaging with the aid of a Brainsight system (Rogue Research Inc). Neuroimaging was performed at the Rochester Center for Brain Imaging, on a Siemens 3T MAGNETOM Trio Tim using 0.5 mm voxels. We confirmed recording locations by listening for characteristic sounds of white and grey matter during recording, which in all cases matched the loci indicated by the Brainsight system.

### Electrophysiological techniques

Single electrodes (Frederick Haer & Co., impedance range 0.8–4 MOhm) were lowered using a microdrive (NAN Instruments) until waveforms of between one and five neuron(s) were isolated. Individual action potentials were isolated on a Plexon system. Neurons were selected for study solely based on the quality of isolation; we never preselected neurons based on task-related response properties. The number of neurons to be collected was determined *a priori* based on exploratory analyses of previously collected datasets and was not adjusted during recording based on analyses performed mid-experiment.

### Eye tracking and reward delivery

Eye position was sampled at 1,000 Hz by an infrared eye-monitoring camera system (SR Research). Stimuli were controlled by a computer running MATLAB (Mathworks) with Psychtoolbox (Brainard and Vision 1997) and Eyelink Toolbox (Cornelissen et al. 2002). A standard solenoid valve controlled the duration of water delivery. The relationship between solenoid open time and water volume was established and confirmed before, during, and after recording.

### Task

*The stop signal task* (**Figure 1A**) is a measure of self-control that provides an alternative approach that avoids some of the limitations of intertemporal choice tasks (Hayden 2016). The task followed standard stop signal paradigm (Logan and Cowan 1984; Logan 1994; Hanes and Schall 1995). Subjects were placed in front of a computer monitor (1920×1080 pixels) with black background. Following a brief (300 msec) central fixation on a white circle (radius 25 pixels, **Figure 1A**), the fixation spot disappeared on the appearance of eccentric saccade target (90px white square, 2.38 degrees, positioned at 288 pixels, 7.62 degrees, in left or 1632 pixels, 43.18 degrees, in right of screen, 50% chance). A *go* trial (67% of stop signal task trials, randomly selected) was indicated by a go cue—a peripheral target, whereas a *stop* trial (33% of trials, randomly selected) was indicated by an additional appearance of a stop signal — a central gray square (90 pixels square, 2.38 degrees) delayed relative to the go cue presentation with a stop signal delay (SSD). On go trials, subjects were rewarded for a saccade towards the go cue and fixating on it for 200 msec; and on stop trials, subjects were rewarded for inhibiting their saccade to go cue, and fixating at the center stop signal for 400 msec. Water rewards were provided as feedback, and they were contingent on subject’s performance. Rewards were always 125 μl. The SSD was initialized to a random floating value between 0 and 0.1 secs for every session (day of recording), and they were adjusted in real time with jumps of 16 msecs by a staircase procedure to stabilize at a delay (SSD-50) with 50% accuracy in the stop trials (Balasubramani et al. 2019). The inter trial interval was 800 msecs (trial baseline).

**Figure 1.**
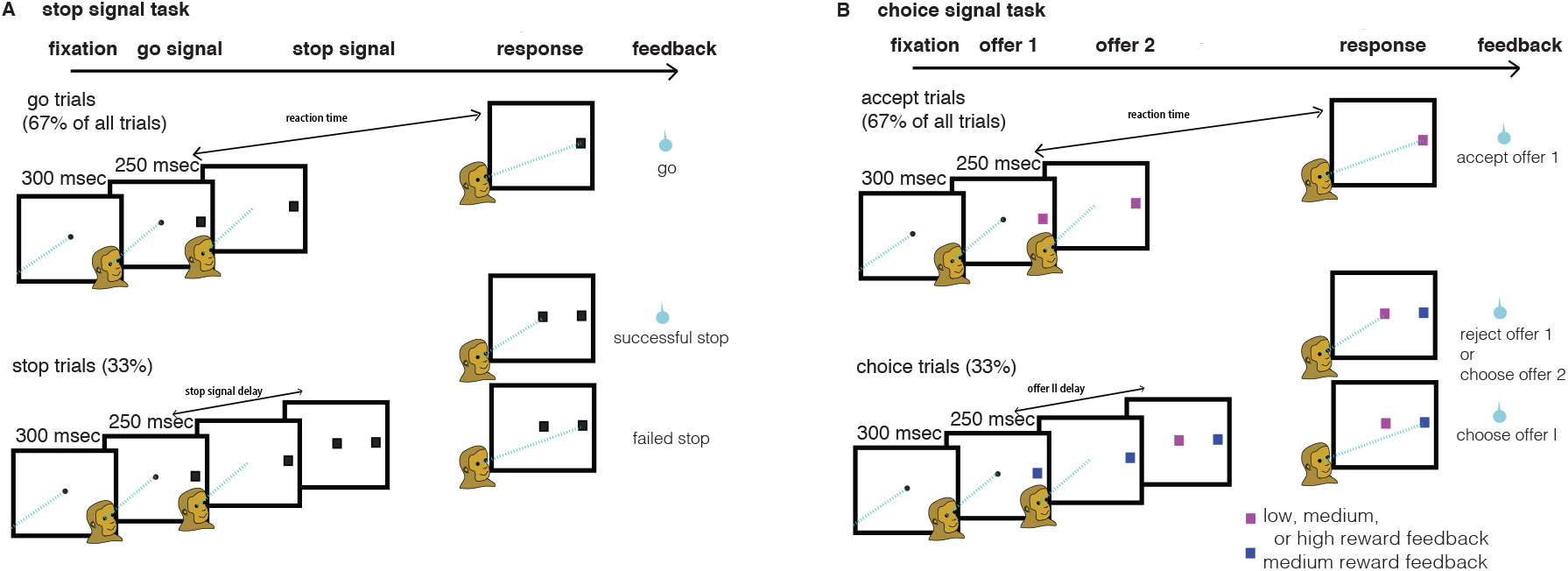
Task framework. (A) stop signal task showing go trials, successful and failed stop trials (B) choice signal task showing accept, choose offer 1 and choose offer 2 (reject) trials.

The *choice signal task* (**Figure 1B**) had a similar task framework to stop signal task. We called the choice signal task’s counterpart of go trials as *accept* trials, and that of the stop trials as *choice* trials. In accept trials (random 67% of the total choice signal task trials), the subjects made a forced choice to a peripheral offer 1 or the first offer (90 pixels square, 2.38 degrees, positioned at 288 pixels in left or 1632 pixels square, 43.18 degrees, in right of the screen, 50% chance). In choice trials (random 33% of the total), an additional second offer (offer 2 or the choice signal) was presented at the center (90 pixels square, 2.38 degrees) delayed with respect to the appearance of offer 1. The choice signal delay for the presentation of the center offer 2 was the same as the current real time SSD followed for the stop signal task in the session. The offers were associated with either low (15μl), medium (125μl), or high (250μl) reward offer sizes, the rewards were respectively indicated by yellow, blue and magenta colored square cues. In accept trials, the low, medium and high offers were presented with equal probability. Similarly for the center choice signal (offer 2) in choice trials, low, medium and high offers were presented in equal probabilities. In contrast, the peripheral offer 1 in the choice trials was always associated with medium reward size (i.e., blue square); this didn’t alter the basic behavioral performance of the subjects in choice trials as a function of value (**Figure 2-1**). So, the cue (blue square, peripheral offer 1) that differentially predicted a choice trial, occurred with probability = 0.6, while probability of a choice trial with non-blue offer 1 was 0.

**Figure 2:**
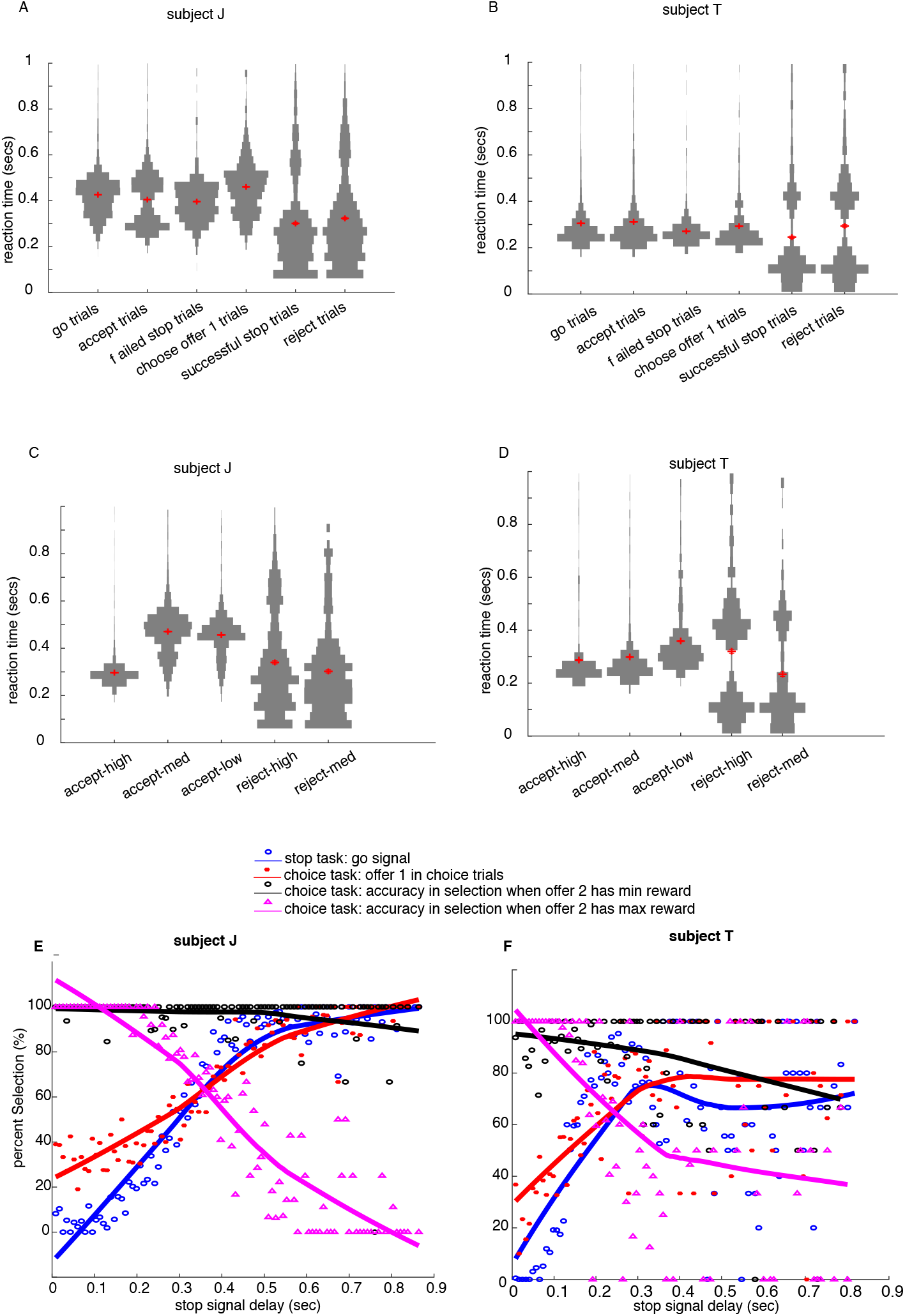
Behavioral results. For subject J, the results are presented in panels (A, C, E) and for subject T in (B, D, F) (A, B) reaction time violin distributions for various trial conditions of stop signal task. The red centroids in the distribution plots denote the mean and the + / -standard error of mean. An extended data representation (**Supplementary material 2)** shows the cumulative histogram as a function of reaction times for various trial types. (C, D) reaction time distributions for various trial conditions of choice signal task. Another extended data (**Supplementary material 3**) shows the session-wise SSD distributions, and SSRT distributions of both subjects, J and T. (E) Inhibition function and accuracy of choices (choosing the best of offers) varied as a function of SSDs. The inhibition function for both the subjects showed sensitivity to the magnitude of stop signal delay (shown in blue lines). (F) The choice functions for different offer reward sizes are shown. The data points in red indicate the percent selection of offer 1 (medium reward size) in choice trials. This varied similar to the inhibition function. The data points in black indicates the accuracy of choice when offer 2 had minimum offer size, and that accuracy was larger compared to another case where offer 2 had maximum reward size (magenta). This suggests that the subjects were relatively more greedy in choosing larger offer sizes (125μl, 150μl) especially when contrasted with an offer with minimum reward size (15μl). The continuous lines in inhibition and choice functions are shape preserving interpolants of the scattered data points obtained by data smoothing with step size of 0.7 secs.

The stop signal task and the choice signal task were stochastically counterbalanced. Specifically, the two tasks were randomly alternated with the constraint that each trail repeated from 1 to 3 trials (number chosen in random).

Subjects had never previously been exposed to decision-making tasks in which counterfactual information was available. Previous training history for these subjects included two types of foraging tasks (see Blanchard et al. 2014; Blanchard and Hayden 2015), several gambling tasks (Farashahi et al., 2018; Yoo and Hayden, 2020; Azab and Hayden 2017 and 2018; Heilbronner and Hayden, 2016), attentional tasks (similar to those in Hayden and Gallant 2013), and two types of reward-based decision tasks (Sleezer and Hayden 2016; Wang and Hayden 2017).

### Behavioral analysis

Reaction time was defined as the time taken to saccade to the peripheral target and was computed relative to the presentation time of the go cue or the first offer 1. In case of a stop trial or a choice trial, the successful cancellation time (or equivalently, the decision time or the choice time in the choice signal task) was the time taken to cancel the saccade by selecting stop signal or the second offer 2, and was computed relative to the presentation of the first offer or the go cue (Schall 1991; Hanes and Schall 1995; Logan et al. 2015).

We used two different methods to compute stop signal reaction time (SSRT). They were 1) median 2) integration methods. We subtracted SSD-50 from the median of go-trial distribution to find stop signal reaction time (SSRT) in the median method. On the other hand, SSD-50 was subtracted from the time point in go-RT distribution whose area was half the distribution total, for computing SSRT in the integration method. We found that SSRT computed through both of the above methods gave nearly equal results, and so, we averaged the values obtained from both methods to obtain final SSRT estimates reported for each subject.

### Statistical methods

Separate PSTH matrices were constructed by aligning spike rasters to the presentation of the go cue or stop signal, appropriately for analysis, for every neuron. Firing rates were calculated in 1 msec bins but were generally analyzed in longer epochs. We normalized by subtracting the mean neuronal firing during inter-trial interval (ITI) time period and z-scored each neuronal data, for bringing them to a common scale of analysis; the normalized data was used for all the decoding analysis mentioned below.

For display, PSTHs were smoothed using 200 msec running boxcars in **Figure 4**.

### Decoding analyses

The decoding analysis used in this study was similar to our recent publication (Balasubramani et al., 2019). Overall, the procedure involved firstly generating of pseudopopulation activation patterns (see Mante et al. 2013; Stokes et al. 2013) similar to that used for a multi-dimensional scaling method for obtaining OFC ensemble activation patterns (for example: Rigotti et al. 2013; Stokes et al. 2013; Cunningham and Byron 2014; Stokes 2015). Second, we performed binary classifications with the OFC activations (for example: Zhang et al. 2018). To generate population activation states as input patterns for the decoding analysis, we first separated all trials of each neuron by trial conditions. Then, we averaged the activity from randomly sampled 10 trials belonging to a condition for a neuron, with replacement, to form activation states. The activation states of all neurons (N = 96) were pooled together to generate a population activation state pattern. 150 unique population activation patterns averaged for some particular time period (either pre-cue or post-cue time periods described below) within the trial were used for decoding analysis.

The shuffling of trials for generating activation states ensured the absence of any temporal relationship between the trials associated with every neuron in the population state ensemble. Hence the generated activation state was independent of any trial-to-trial temporal relationship between neurons, that were representative of “simultaneous” recording. Therefore, it is valid for our method to be applied for ensemble decoding irrespective of its simultaneous recording nature, i.e., both for simultaneous and non-simultaneous recordings.

The network had a single hidden layer with 100 hidden nodes, and 2 output nodes each representing one target condition for classification. The number of input nodes equal to the total number of neurons used for analysis = 96 (from two subjects). The network weights were initialized to small random numbers between −0.01 and 0.01.

The *back-propagation algorithm* was used for training the decoders (Werbos 1974; Rumelhart et al. 1986; Rumelhart et al. 1988; Haykin and Network 2004). In the below eqn. 1, the input nodes are denoted by subscript, *k*, hidden nodes by subscript, *j*, and output nodes by subscript, *i*. Output error, *e*, associated with the network’s response for the *p*’th input pattern is then given by

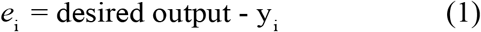

where *y*_i_ is the *i*’th output node response, and desired output is 1 / 0 if the *i*’th output node is associated with target trial condition for the corresponding input pattern (e.g., successful stopping, failed stopping). Total output error over all input patterns is computed by,

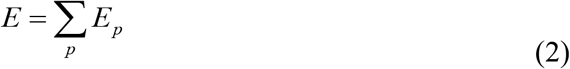

Network’s objective was to minimize the squared output error for the *p*’th pattern as denoted by eqn. (3).
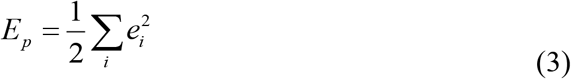

A hyperbolic tangent function (*g*) with slope = 5 was used as the activation function for every node in the network. The learning rate was set to 0.001 for the network. The node with the maximal activation value in the output layer was taken as the network output response.

We focused our analyses on two key time periods of the trial (1) the 300 ms epoch before the trial started called the pre-cue epoch, which was actually the fixation time before the presentation of any stimulus targets. (2) The variable time after the go cue (or the first offer in case of the choice signal task) and before the reaction timepoint called the post-cue epoch.

For pre-cue decoder, the training data were population activation states generated by averaging neural activity from the pre-cue epoch of a trial respectively. Similarly, the training data for the post-cue decoder was computed by averaging the post-cue epoch. As mentioned earlier, the pre-cue and post-cue decoders were two class decoders with two output nodes, that distinguished two different trial types. We used the decoders to identify various trial types such as successful vs. failed stop trials, go vs. successful stop trials in the stop signal task. In choice signal task, the accept trials with were divided based on non-high (containing the low and medium offer sizes) vs. high offer sizes to classify into two-classes. Accept vs. reject trial types were also analysed as two different classes in the decoders.

The artificial neural network was run for *n* = 100 instances with different random weight initializations for obtaining average output performance. Training procedure in all instances converged to classification accuracy of above 80%, and the converged weights at the end of training were used for the testing phase of the decoders. The epoch used for generating activation states for training were from one task and for testing was activation states of the same epoch but from the alternate task. The testing was done on an independent test dataset that was distinct from the one used for fitting. Specifically, we did a 60% training / 40% testing split of our population data, and we made sure different trials were used for generating the population training and testing states. We also ensured each training and testing set was stratified-it had equal proportion of different trial types (classes) during training and also testing. We verified whether the decoding performance was significantly higher than 50% using chi-square test (p < 0.05). Furthermore, we performed permutation tests as a control. In the permutation tests, we shuffled the labels for train and test 5K times for generating a null distribution, and we found its 95 percentile value was 80 percent decoding performance. To that end, we present only the significant results with classification percentages greater than 80 (95 percentile null distribution value) in **Table 1**.

**Table 1:** overlapping functional ensemble codes for stopping and choice: Decoding overlapping neural codes between stopping and choice tasks. The rows indicate the training while the columns indicate the testing signals. The different highlighting colors represent the differences in the task signals used as training and testing the decoders. All the decoders were trained well with greater than 80% performance, and the testing results survived the statistical corrections (chi-square test, p<0.05) and the performance results were well above 95 percentile threshold of permutation test (=80% performance)

| TRAINING \ TESTING |  |  | % decoding of Pre-cue epoch |  |  |  | % decoding of Post-cue epoch |  |  |  |
| --- | --- | --- | --- | --- | --- | --- | --- | --- | --- | --- |
|  |  |  | Stop signal task |  | Choice signal task |  | Stop signal task |  | Choice signal task |  |
|  |  |  | Failed vs successful stop | Go vs failed stop | Accept vs. reject | High vs non-high accept | Failed vs successful stop | Go vs failed stop | Accept vs. reject | High vs non-high accept |
| Pre-cue epoch | Stop signal task | Failed vs successful stop |  |  | 85.2 | 87.2 |  |  |  |  |
|  |  | Go vs failed stop |  |  | 92.4 | 90.1 |  |  |  |  |
|  | Choice signal task | Accept vs. reject | 98.7 | 97 |  |  |  |  |  |  |
|  |  | High vs non-high accept | 99.3 | 95.7 |  |  |  |  |  |  |
| Post-cue epoch | Stop signal task | Failed vs successful stop |  |  |  |  |  |  | 92.4 | 88.8 |
|  |  | Go vs failed stop |  |  |  |  |  |  | 94.4 | 84.5 |
|  | Choice signal task | Accept vs. reject |  |  |  |  | 99 | 94.4 |  |  |
|  |  | High vs non-high accept |  |  |  |  | 97.7 | 95.7 |  |  |

trained in a choice signal task but tested in stop signal task
trained in a stop signal task but tested in choice signal task

The decoder was sensitive to the temporal activations of each single neuron independently in an ensemble. Our decoder worked on the activations of single neurons, in an independent fashion, and related the ensemble activation state to behavior. That is, the decoder cared about the temporal activity within a neuron i.e., the event locked response activity of every independent single unit. Specifically, we had suggested that the decoding efficiency to be higher for an ensemble activity for precise time periods from the presentation of the stop signal in our recent study (Balasubramani et al., 2019).

This approach of using a multidimensional state decoding of an ensemble with “non-simultaneously” recorded cells was used by multiple other studies (example: Gochin et al. 1994; Thomas et al. 2001; Averbeck et al. 2006). This is a subtler point, but important. Our method is logically equivalent, in many ways, to several other dimensionality reduction approaches that were used for neuronal ensembles (for a review, see (Cunningham and Byron, 2014), and other studies, (Rigotti et al., 2013; Stokes, 2015; Stokes et al., 2013). It is well known that these methods can be done with no problems on asynchronously collected cells, for the same reason that our similar methods can. We had also applied our methods to simultaneously recorded smaller ensembles and found consistent results (Balasubramani et al., 2019)- so our decoding method is general enough to analyse the activation states of neural ensembles either recorded simultaneously or constructed as pseudo-population (Averbeck et al., 2006; Gochin et al., 1994; Thomas et al., 2001).

## RESULTS

### Behavior in the stop signal task and choice signal task

Subjects performed two interleaved tasks, a classic ***stop signal task*** (Hanes and Schall, 1995; **Figures 1A** and **Methods**), and a novel ***choice signal task***. We reported some of the results on the stop signal task previously (Balasubramani et al, 2019). Our focus here is on comparing those stop-signal results with results in a second task (the choice signal task) that was collected in interleaved trials. Those data were not previously analyzed.

The ***stop signal task*** consists of two kinds of randomly interleaved trials: *go* trials (67%) and *stop* trials (33%). On either of these two trial types, following a central fixation, subjects first saw an eccentric target (*go cue*). On *go trials*, a juice reward was provided following a successful saccade to the cue. On *stop trials*, a second signal (*stop signal*) appeared at central fixation point, and the subjects had to countermand the previously instructed saccade for a reward. Failure to inhibit resulted in no reward. These were called *failed stop trials* while successful inhibition resulted in a reward and were called *successful stop trials*. The delay between the go cue and the stop signal is called the stop signal delay (SSD); we used staircase procedure in real time during each session or the day of recording to stabilize at a delay with 50% success in stopping (SSD-50, see **Methods**), and computed the stop signal reaction time (SSRT, Logan and Cowan 1984; Logan 1994; Verbruggen and Logan 2008). The SSD-50 was 0.27 sec for subject J and 0.15 sec for subject T and SSRT was 0.14 sec for subject J, and 0.12 sec for subject T (Balasubramani et al, 2019).

The randomly interleaved ***choice signal task*** was designed to have a structure isomorphic to the stop signal task. It used interleaved *accept trials* (66% of trials), analogous to go trials, in which the subject made a forced choice by accepting a first offer presented peripherally on the screen (offer 1), and *choice trials*, analogous to the stop trials, where the subjects were presented with not just the peripheral offer but also a second offer (offer 2) at the center. The trials in which the subject chose the offer 1 were called *choose offer 1* trials; trials in which the subject chose center offer 2 were called the *reject trials* or *choose offer 2* trials. Reject trials were analogous to successful stop trials (see **Methods** for details; the second offer was presented 33% of times as a choice signal with the same delay as the current real time SSD of the stop signal task). In the accept trials, the offers were either low (yellow color cue), medium (blue), or high (magenta) in reward size value (**Figure 1B**), each presented with equal probability; And in the choice trials, the peripheral offer was always with medium reward size (blue) while the center one was either low, medium or high in offer size, presented with one thirds of probability each. Subjects learned this task well, and successfully chose offers with relatively high reward about 89% of the times (**Supplementary material 1**).

Reaction time patterns in the two tasks were similar. Reaction time distribution metrics between the stop signal task and the choice signal task for both the subjects J and T appear in **Figure 2**. Reaction times on accept trials (choice signal task) correlated with the go trial reaction times (stop signal task) significantly across sessions (subject J: Pearson correlation, r = 0.90, p <0.001; subject T: r = 0.86, p < 0.001). That means day to day variability in performance affected the two tasks in a similar way. Likewise, reaction times on *choose offer 1* trials were correlated with those on the failed stop trials (subject J: Pearson correlation, r = 0.90, p < 0.001; subject T: r = 0.72, p = 0.005). Choice accuracy (selecting an offer with relatively larger reward size) when found with respect to the delay in the presentation of offer 2 (i.e., SSD) significantly correlated between subjects, particularly, as the delay between the presentations of offer 1 and offer 2 increased, the accuracy in choosing the best of the offers decreased (**Figures 2E** and **2F**; **Supplementary material 1**). A similar behavior was seen in the stop signal task, where there was a decrease in the percentage of successful saccade inhibition (i.e., inhibition accuracy) with increase in SSD (**Figures 2E** and **2F**). Note that subject T had higher variance in his inhibition function during longer delays but the behavior was still monotonic, allowing us to study his neural and behavioral data as a function of delay time, particularly around the SSD-50 and SSRT periods (**Figure 2F**); these longer delay trials did not affect the interpretation and results of our study, and our neural decoding analyses described in this study do not focus on the effects of SSD on behavior.

The similarity in the trial-by-trial effects between the tasks provides tentative support for the hypothesis that subjects treat these tasks in analogous ways (**Figure 3**). Further supporting this idea, the reaction times of the go trials that followed a successful stop trial were correlated with the reaction times of the accept trials that followed a reject trial in the choice signal task (Pearson correlation, r = 0.71, p < 0.001). Likewise, we saw a significant correlation between the reaction times of saccade trials that followed another saccade trial in both the tasks (go trials / accept trials; r = 0.88, p < 0.001).

**Figure 3:**
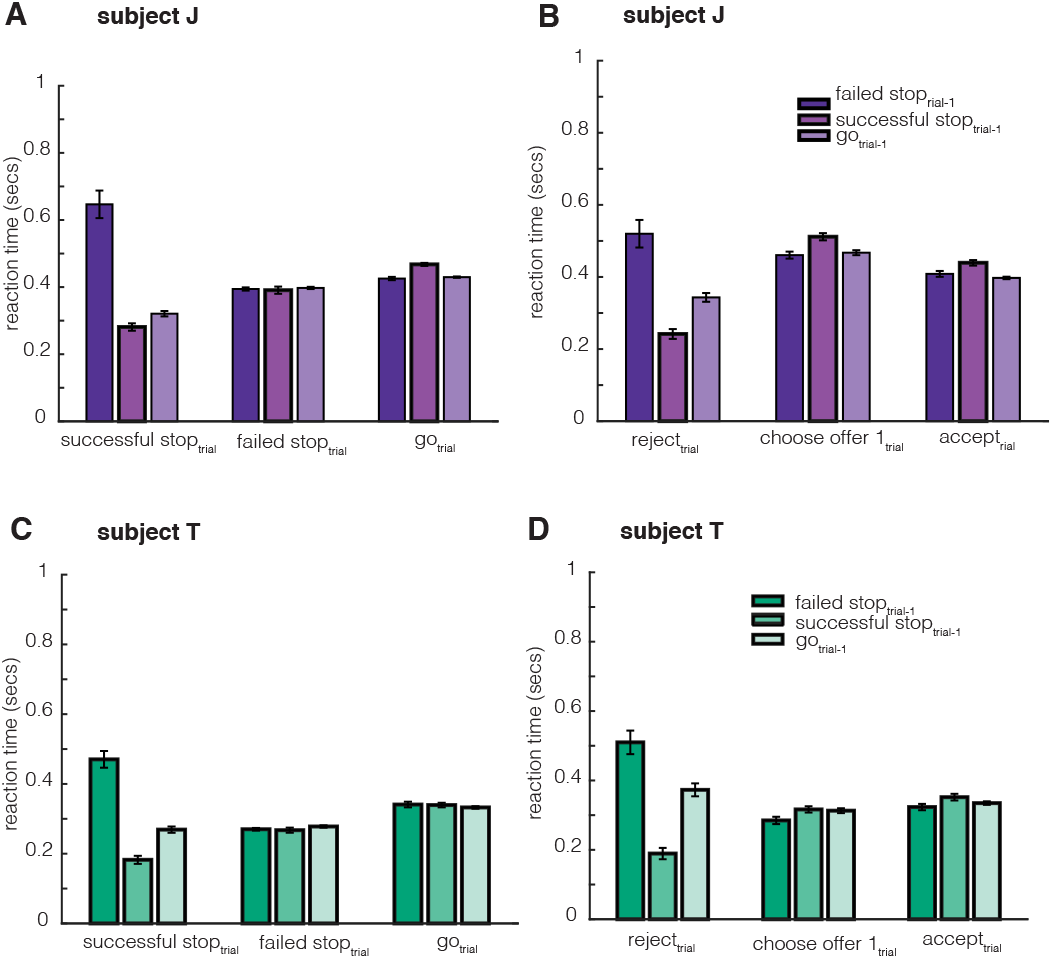
Trial-to-trial behavior effects. For various trial conditions of stop and choice signal ask-we found trial history based effects in reaction time behavior in (A, C) stop signal task and (B, D) choice signal task. Error bars represent SEM. Accept trials were generally longer when followed an reject trials in contrast to following another accept trial (subject J figure 3B: 0.05 s, tstat = 3.41, p < 0.001; subject T figure 3D: tstat = 2.06, p = 0.039). Similar effects were found for stop signal task: Successful stop trials were shorter when after another successful stop in contrast to following a failed stop trial (subject J figure 3A: 0.36 s, t-stat = 11.33, p < 0.001; subject T figure 3C: 0.29 s, t-stat = 11.88, p < 0.001).

### Stopping and choice at the level of single neurons

We recorded responses of 96 neurons (52 in subject J and 44 in subject T) in Area 13 of the OFC (see Balasubramani et al, 2019). Note that while this number of neurons is smaller than in some other studies, it is sufficient to have detected the effects we discuss and thus suits our purposes. We show responses of three example neurons in **Figure 4**. As is typical in OFC, neurons had diverse tuning (coding) profiles.

**Figure 4:**
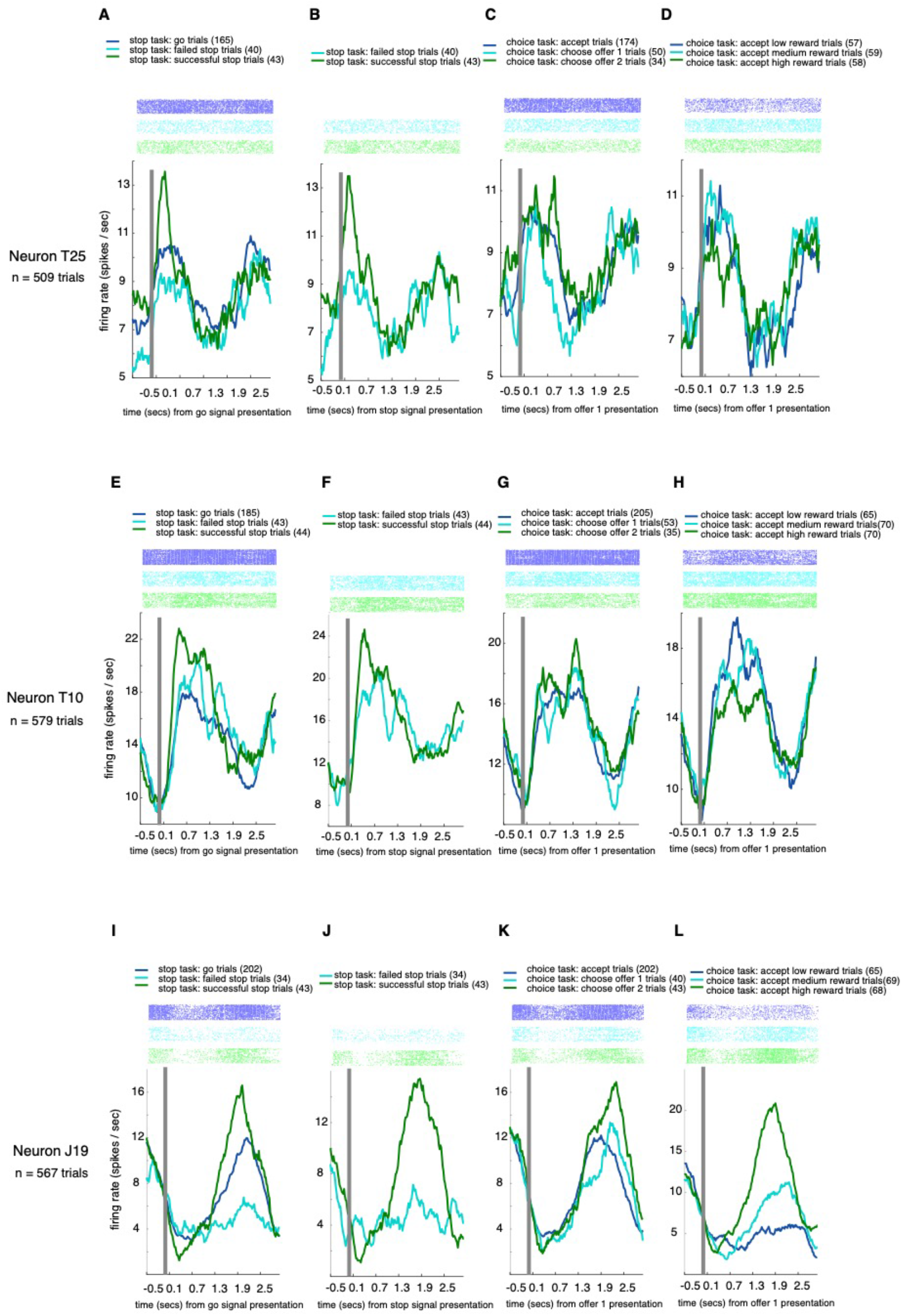
Illustration of sample neurons. We present neuron T25 in panels A-D, neuron T10 in panels E-H, and neuron J19 in panel I-L. The first panel of every neuron shows the activity in stop signal task when aligned to the go cue (A, E, I), and that when aligned to the stop signal in the second panel (B, F, J). When aligned to the go signal, we show the go, successful and failed stop trials’ mean neural activity. And when aligned to the stop signal, we present the mean neural activities for the successful and failed stop trials. The last two panels for every neuron show the matching trials from choice signal task. The panels (C, G, K) present the accept, choose offer 1 and choose offer 2 (reject) trials of each neuron, while the panels (D, H, L) present the accept trials of choice signal task with different offer sizes (high, medium and low offer sizes) of each neuron. Some significant differences in neural firing rates between trial types were seen in the stop signal task: Neuron T25 differentiated successful and failed stop trials in pre-go cue and post-go cue time periods (Figure 4A; pre-cue: ttest, p = 0.01, post-cue: p = 0.02). The inhibition code could be found even with respect to the presentation of the stop signal, the neuron differentiated the successful and failed stop trials in the post-stop signal time period before the stop signal reaction time (p = 0.01, Figure 4B). Another neuron, T10, differentiated the successful and failed stop trials in the post-go cue epoch, (ttest, p = 0.04, Figure 4E) and similarly, it also differentiated go vs. successful stop trials (p = 0.002, Figure 4E) in the same post-go cue epoch. Neuron J19 differentiated successful and failed stop trials even before stop signal reaction time in the post-go cue epoch (t-test, p = 0.05, Figure 4I), as well as in the post-stop signal time period before stop signal reaction time (p = 0.019, figure 4J). In the choice signal task based trials (Figure 4C), the neuron T25 differentiated choose offer 1 and choose offer 2 (reject) trials in the post-offer 2 presentation time period before SSRT (p = 0.04, figure 4C). The neuron T10’s and J19’s mean firing activity in the accept trials were negatively correlated to the size of offers during feedback (Pearson correlation, r = −0.16, p = 0.017 for T10; r = 0.40, p < 0.001 for J19).

We used responses from all of the 96 neurons for the following analyses. We focused our analyses on comparing stopping and choice processes only in two key time periods of the trial: (1)*pre-cue epoch:* the 300 ms epoch *before the trial* started, which is related to the fixation time before the appearance of any stimulus targets and that can inform about proactive signaling. (2) *post-cue epoch*: the variable time after the go signal (first offer 1 in choice signal task) but before the reaction time that can inform about reactive signaling.

For the choice signal task, 45.83% of cells showed positive trial related firing response compared to the baseline, and 51.04% showed negative trial related response, and the rest weren’t significantly modulated (t-test on firing rates in post-cue epoch, α=0.05). We saw a similar pattern in the stop signal task: In 44.79% of cells, the average firing was higher during the trial compared to baseline and 53.12% had responded with a negative average firing to the baseline, the rest (2.09%) showed no change.

### Distinct neural activities for actions and rewards in the two tasks

Our key question is how firing rate patterns in the two tasks relate to each other. Focusing on the post-cue epoch, and we first asked whether the patterns for go / stop actions in the stop signal task have any similarity to the accept / reject choice patterns. In other words, does *going* in the former task correspond – neutrally - to *accepting* in the latter task? This pattern is expected because the actions are matched. We regressed the firing rate against the action variable (saccade or inhibition of saccade) in the two trial types; coefficients were significantly correlated (r = 0.28, p < 0.001). This, positive correlation indicates the neurons encode action information in a similar fashion in the same direction. We also correlated unsigned coefficients; a positive correlation is evidence that the two relevant neural populations overlap more than chance (Azab and Hayden, 2017). Here we found a positive correlation as well, indicating that the same groups of neurons are involved in the two tasks (r = 0.65, p < 0.001). Neither result is particularly surprising, although they provide valuable confirmatory evidence and set the stage for our subsequent results.

Next, we asked how neural patterns for the two tasks relate in terms of reward magnitudes associated with chosen options (also called the value of chosen option). In the stop signal task when the stop signal is presented, only successful stop trials provided rewards, that means the action and reward magnitude identifiers were one and the same in stop trials and they cannot be used differentially identify reward magnitude patterns. Moreover, because of this confound, any similar effects between stop signal task and choice signal task patterns may reflect their shared actions (and OFC has spatial tuning, Feierstein et al. 2006; Roesch et al. 2006; Strait et al. 2016; Yoo et al. 2018) instead of chosen option values. In order to understand the reward value effects, we controlled for the trials in the choice signal task. We used only accept trials, thus the action was the same – a saccade, in the choice signal task. We regressed the neural activity in the post-cue epoch and the actions (saccade or inhibition of saccade) in the stop trials, and, in parallel, we regressed the post-cue neural activity with the received reward sizes only in the accept trials. We then asked whether the regression coefficients from two tasks were significantly related. If they were related, it would imply OFC had similar codes for action value and reward value. Interestingly, the regression coefficients computed for each neuron in two different tasks weren’t related for the post-cue epoch (Pearson correlation, r = 0.06, p = 0.58). Moreover, the unsigned coefficients weren’t correlated (Pearson correlation, r = 0.10, p = 0.31). This implies OFC neuronal codes for stopping and choice value don’t have a simple linear relationship in the post-cue epoch.

In a previous study, we had found that subjects perform reward value comparisons between offers for making their choice (Strait et al., 2014). Since go / stop is associated with a distinct outcome in stop trials, we next asked whether the principles regulating choice can be applied to stopping, that is performing reward value comparison between actions. To answer this question, in the choice signal task, we estimated how the neural activity during choice was related to reward value differences between offers. Specifically, we regressed the neural firing in the post-cue epoch of choice trials with their offer size differences (offer 2 – offer 1). And in the stop signal task, to estimate how the neural responses were related to actions, we regressed the firing activity in the post-cue epoch of the stop trials with the action regressor (saccade or inhibition of saccade). We compared the computed coefficients for each neuron, and we didn’t find any significant relationship (Pearson correlation between the signed coefficients: r = 0, p=0.95), but of significance was the relationship between the unsigned coefficients (r = 0.52, p<0.001). This result with the unsigned coefficients suggests that an overlapping subset of neurons assists both choice and stopping in the OFC, although their codes aren’t relatable in a simple and linear fashion as implied by the signed coefficients.

### Overlapping functional ensemble codes for stopping and choice

In the earlier section, we saw action related and reward related relationships at the level of firing rates between the tasks. Firing rates for actions were highly correlated between tasks, and these codes weren’t simply related to the reward signals. We then wanted to examine the extent of overlap between the tasks at the level of population ensembles. Notably, population averages revealed no significant differences between stopping / going trial types in the stop signal task as well as reject / accept trial types in choice signal task (**Figure 5)**.

**Figure 5:**
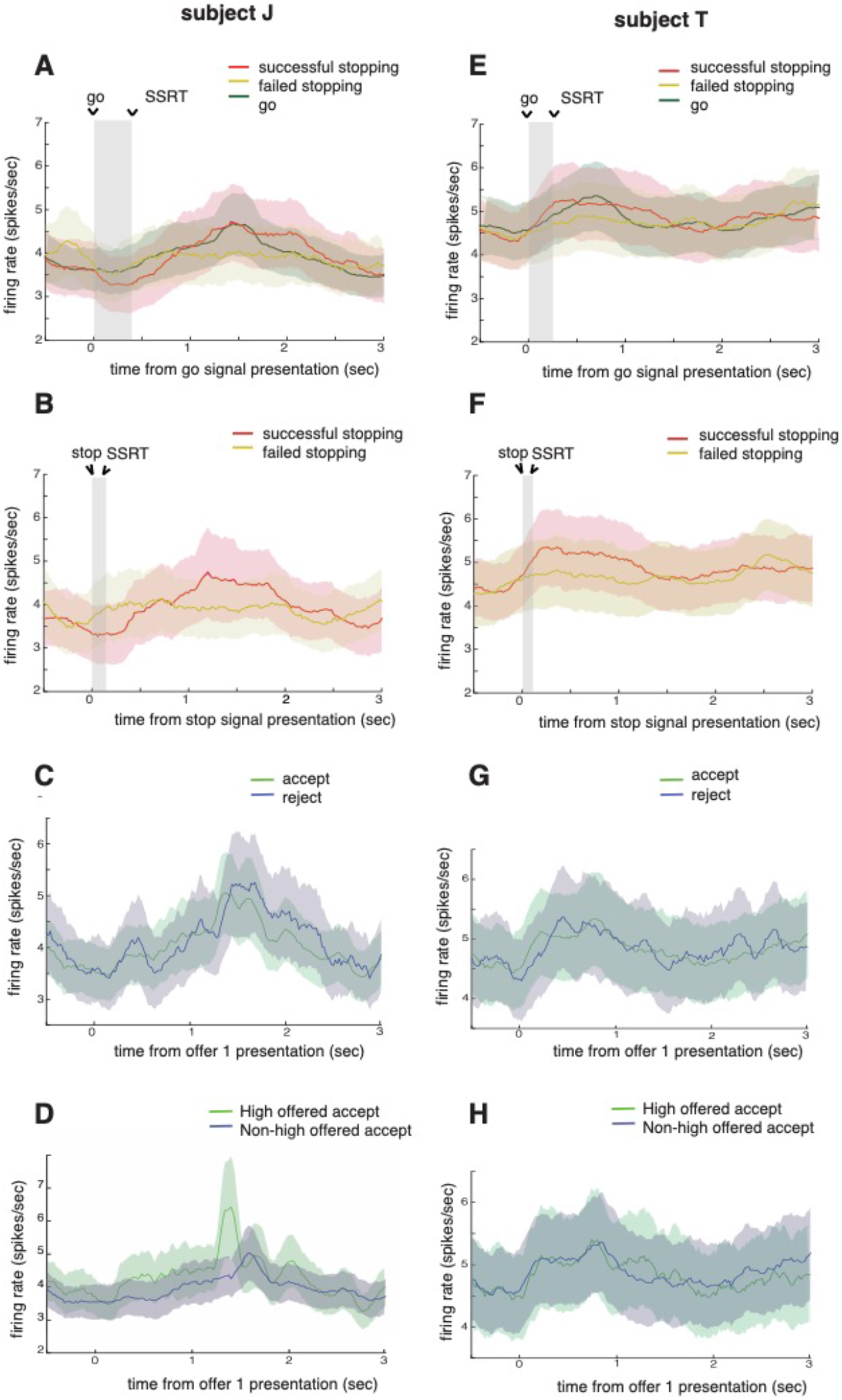
population averages provide weak information about stopping and rewards. Population activity for successful stopping and failed stopping with respect to **(A, E)** go cue presentation and **(B, F)** stop signal presentation, for subjects J and T. Time from start of the go cue to SSRT is shaded in panels A and E (and stop signal presentation to SSRT in B and F). Data for all SSDs are averaged to present the signals during successful and failed stop trials. They didn’t reveal significant information about the pattern of stopping. In the same lines, population responses for accept and reject trials for choice task in subjects J and T, respectively, are shown in **(C, G);** they didn’t reveal significant information about choice patterns. Finally, the population averages for high and non-high offering accept trials are presented in **(D, H)** for subjects J and T, respectively. Error bars in all panels denote SEM.

There is some evidence to suggest that ensemble patterns can code for neural information that may not be conveyed at the level of single units (Zemel et al. 1998; Averbeck et al. 2006b; Meyers et al. 2008; Morcos et al. 2018). We therefore made use of neural network decoders, which provide the most direct way to detect patterns in the ensemble with minimum number of assumptions. This approach has been used successfully in several other studies (for example: Rigotti et al. 2013; Stokes et al. 2013; Cunningham and Byron 2014; Stokes 2015). We trained two different decoders, first one was trained on stop signal task and tested on choice signal task; the second one was the reverse (and was used for confirmation). These analyses used the same two epochs as above (**Table 1**). In any instance, population activation patterns (N=96) were generated (see **Methods**) from only one of the epochs (pre-cue, post-cue) for both training and testing, and they were fed in as input to the decoder. In case of shared neural patterns underlying stopping and choice, a decoder trained on just one of tasks would perform with high efficiency when tested on the other task (the results should survive statistical corrections for significance: chi-square test, p < 0.05, ≥ 95 percentile threshold in permutation test, see **Methods**). Our results below indeed suggest the existence of overlapping ensemble patterns between stopping and choice.

We examined the decoding performance of some key patterns or trial differences in stopping and choice: (1) successful vs. failed stop trials, where the trials not just differed in actions (only successful stop trials were associated with inhibition of saccade) but also differed in reward sizes (no reward during failed stop). (2) go trials vs. failed stop trials, where the trials only differed in reward sizes (only go trials were associated with reward) but not in actions (saccade). Their counterparts in choice trials: (3) accept vs reject trials, where the trials differed both in actions (only reject trials were associated with inhibition of saccade) and in offer values. (4) high vs. non-high offering accept trials of the choice signal task, where the trials only differed in reward sizes but not in actions (saccade). The results (**Table 1**) show that the decoders trained to distinguish the trials belonging to one task but tested on the trials belonging to the other, performed with greater than 80% efficiency, indicating strong overlap between stopping and choice ensemble codes, both in the pre-cue epoch and post-cue epoch. Particularly, the decoders trained on choice signal task activations had over 97% efficiency in decoding the successful vs failed trials of the stop signal task (**Table 1**, highlighted in yellow), suggesting a relatively stronger overlap of stopping to choice than the vice versa.

Reward promotes choice of the option it is associated with. The results show that OFC ensembles make use of similar patterns for stopping and offer value (Hayden 2018), and in terms of direction of the effect, ensemble patterns for stopping are more similar to choice patterns. Similarities between neural patterns of both the tasks may promote the idea of shared strategies between stopping and choice related decisions.

## DISCUSSION

We examined responses of neurons in area 13 of the OFC in two tasks, one an implementation of a classic stop signal task and the other a simple choice signal task with similar structure. Although the economic role of this region in choice is well established, its role in stopping is not. Our earlier finding that OFC ensembles predict inhibition both before the trial and immediately before the stopping response suggest that this region does participate in regulation of stopping (Balasubramani et al., 2019). The timing of the two stopping-related patterns is reminiscent of the times associated with proactive and reactive control, respectively (Hanes et al. 1998; Stuphorn et al. 2000; Ito et al. 2003; Braver et al. 2007; Chikazoe et al. 2009; Chen et al. 2010; Stuphorn et al. 2010; Braver 2012; Stuphorn and Emeric 2012; Majid et al. 2013). In the present study, by interleaving the tasks for stopping and choice, we show, first, that it is largely the same neurons participating in both processes. Second, through population analysis and pattern decoders, we show the ensemble patterns that differentiate value of offers underlying the choice signal task can distinguish failed from successful stopping. These results thus support the hypothesis that stopping and choice signal reflect common computations occurring in overlapping circuits.

There is some reason to think that stopping and choice may spring from shared processes. For example, several psychiatric conditions, including depression and addiction, impair both processes, and greater impairment of both is associated with greater disease progression (Nestler et al. 2002; Iacono et al. 2008; Volkow et al. 2011). Second, both are closely associated with, and empirically linked to, the broader concept of self-control (Inzlicht et al. 2014; Berkman et al. 2016; Shenhav 2017). Earlier studies have referred to both inhibitory control and choice processes to reflect upon the nature of “self-control”. Some examples include controlling of choice towards a tempting sub-optimal option (e.g. Baumeister and Newman 1994; Mischel et al. 1996; Muraven and Baumeister 2000; Shenhav et al. 2013; Shenhav 2017), and it extends to even avoiding the sub-optimal ones (Fujita 2011; Berkman et al. 2016). One theory of self-control sees it as a result of competition between two systems-impulsive (hot) and controlled (cold) systems (James 1890; Baumeister and Heatherton 1996; Carver 2005) this relates to the view of inhibitory control (that studies the ability to successfully inhibit a response process) as a form of self-control—where a cold system tries to win over the hot one. On the other hand, many studies suggest the view of self-control as an economic decision, i.e., a comparison of two different utility options (McGuire and Kable 2015; Berkman et al. 2016).

One of the major motivations for this study was to test a set of hypotheses that have emerged from our laboratory in recent years (Hayden 2018; Hayden and Moreno-Bote 2018; Yoo and Hayden, 2018). These ideas, which we sometimes call the foraging view of economic choice, are inspired by, and somewhat similar to, earlier models of choice, especially work of Krajbich and colleagues and Kacelnik and colleagues (Krajbich et al. 2010; Kacelnik et al. 2011). In foraging theory, decisions are generally framed as accept-reject (Stephens and Krebs 1986; Stephens and Anderson 2001; Kacelnik et al. 2011; Blanchard and Hayden 2015). In our view, attention focuses on one option at a time, the subject evaluates the attended option relative to the value of rejection, and then the subject either accepts or rejects it. In a two-option choice, the first attended option has a high value of rejection, since the cost to inspect the second is very low, and rejecting the first does not preclude its choice a few hundred milliseconds later. In this view, comparison is, generally speaking, an emergent process. That is, attending to each option triggers a slow evaluation process, one whose speed is determined in part of the value (or strength of evidence) of the attended option (Krajbich et al. 2010). The comparison occurs in a de facto manner, because the stronger option is more likely to reach the threshold sooner. In practice, this approach has many similarities with traditional accept-reject decisions. However, the key differences include asynchronous consideration of options and a change in how choice occurs in single and multiple (more than two) option cases. Specifically, an important difference comes from the fact that choice process involves steps in which the subject chooses between selecting and rejecting the attended option – between going and stopping. Our results provide neural evidence consistent with this idea.

These results obtained from our overlapping task design provides evidence in favor of the hypothesis that the neural processes that regulate stopping overlap with the processes that regulate economic choice. One limitation of network decoders used in the study is they do not provide any ready insight into the mechanisms of how neuronal outputs are combined to drive action. Doing so is not our goal here. Instead, our main goal is to understand how neural patterns relate across the two categories of tasks.

Overall, our results suggest one core function of OFC may be to generate an abstract regulatory signal to feed into a cascade of downstream structures that ultimately determine choice (Hunt and Hayden 2017; Balasubramani et al. 2019; Yoo and Hayden, 2018). In this way, it may be similar to other regions, especially cingulate cortex but also striatal regions (Hillman and Bilkey 2010; Shenhav et al. 2013; Sleezer et al., 2017; Sleezer et al., 2016). In the context of economic choice, this signal will resemble a value signal; in other cases, it will correlate with other relevant task variables. This view is consistent with the idea that choice and control processes both reflect a gradual transformation of network processes across brain regions obeying principles of distributed systems (Hunt and Hayden 2017; Eisenreich et al. 2017; Balasubramani et al. 2018). One benefit of view is that it provides a basis for the observed role of OFC and their adjacent structures in self-control (McClure et al. 2004; Kable and Glimcher 2007).

## EXTENDED DATA

### Supplementary material 1

Supplementary material 1:
| Trial type |  | Reaction time distribution medians |  |
| --- | --- | --- | --- |
|  |  | (s) |  |
|  |  | *subject J* | *subject T* |
| Stop signal task | Go trials | $0.43 \pm 0.09$ | $0.27 \pm 0.12$ |
| | Failed stop trials | $0.39 \pm 0.09$ | $0.25 \pm 0.08$ |
| | Successful stop trials | $0.25 \pm 0.20$ | $0.15 \pm 0.22$ |
| Choice signal task | Accept trials | $0.41 \pm 0.12$ | $0.27 \pm 0.13$ |
| | Choose offer 1 trials | $0.46 \pm 0.12$ | $0.25 \pm 0.10$ |
| | Choose offer 2 (reject) trials | $0.28 \pm 0.19$ | $0.20 \pm 0.23$ |

| Choice signal task |  | *subject J* | *subject T* |
| --- | --- | --- | --- |
| Variables |  |  |  |
| Reaction times versus value of offers | Forced choice trials- offer 1 | Pearson correlation, $r = 0.52$, $p < 0.001$ | $r = 0.43$, $p < 0.001$ |
| | Choice trials – offer 2 | $r = 0.29$, $p = 0.003$ | $r = 0.27$, $p < 0.001$ |
| Preference for offer 1 than offer 2 | Choice trials | 57.6%<br>(chi-square test, chi-square = 5.78, p = 0.016) | 56.25%<br>(chi-square stat = 3.38, p = 0.066) |
| Preference for offer 1 versus stop signal delay | Choice trials | slope of psychometric sigmoidal fit = 5.53 | Slope = 6.93 |
| Accuracy (choosing best offer) | Choice trials | 89.25% | 89.8% |
| Accuracy versus stop signal delay | Choice trials | slope of psychometric sigmoidal fit = -4.93 | Slope = -5.3 |
| Neuronal coding for rewards | Offer 1 | 21.15% | 9.09% |
|  | Offer 2 | 11.5% | 9.09% |

### Supplementary material 2

The reaction time distributions are accumulated in a cumulative plot to show the differences between trial types for both tasks in subjects J and T.

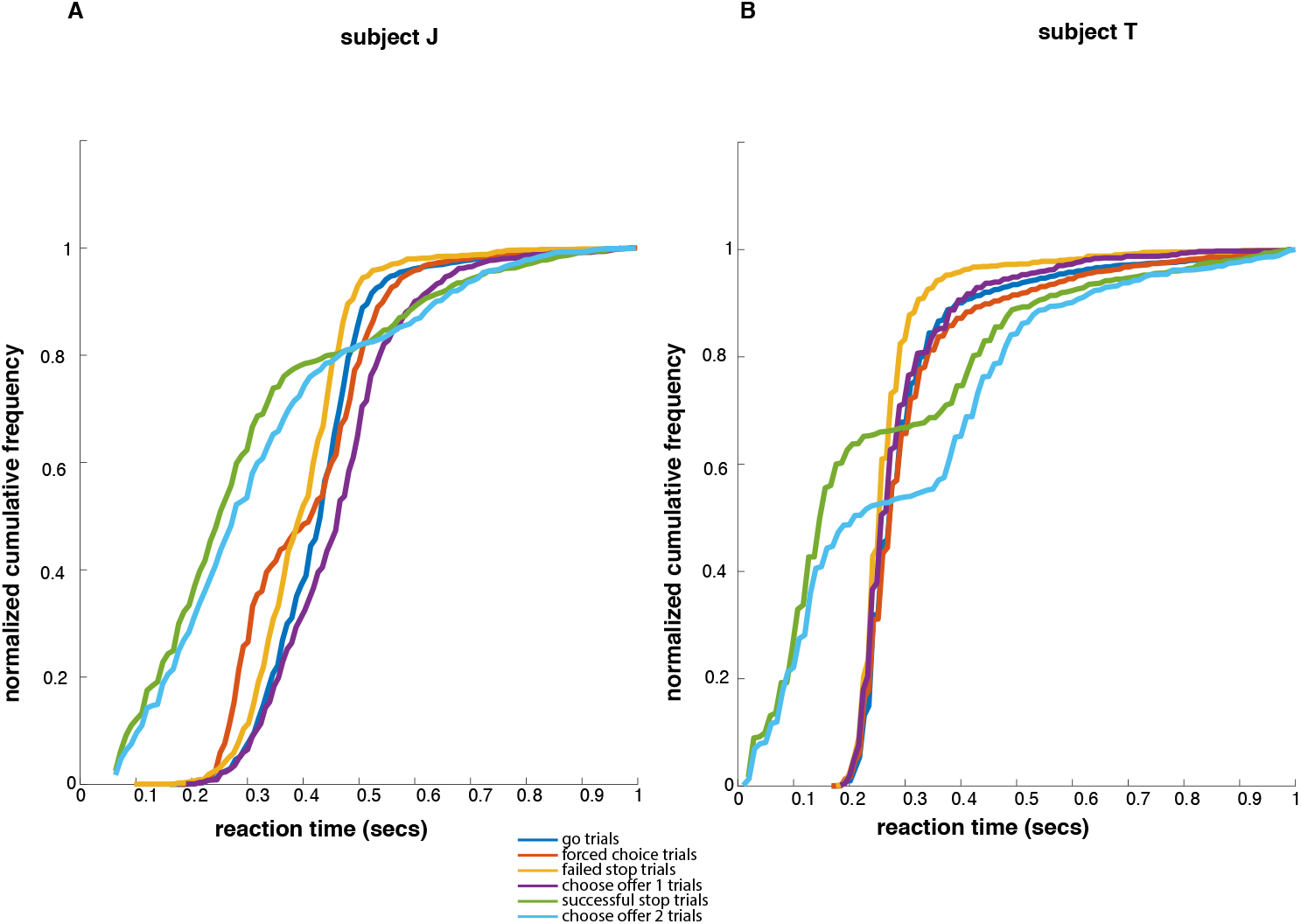

### Supplementary material 3

We collected our data on day basis and it was for 19 days in subject J and 13 days in subject T.

The SSD starting point is a random floating value between 0 and 0.1 secs. Below in the top subplot of the figures, we show SSD distribution plots for each session in both subjects. In the bottom subplot, we show that a representative session’s SSD (current value) trail oscillate around a mean SSD value (SSD-50, as evident from the triangular curves) through trials, both in subject J and T, suggesting they stabilize and converge. The starting point of the trail is when the neural recording started for the day. (Note that the start of neural recording isn’t exactly the start of the task session for the day).

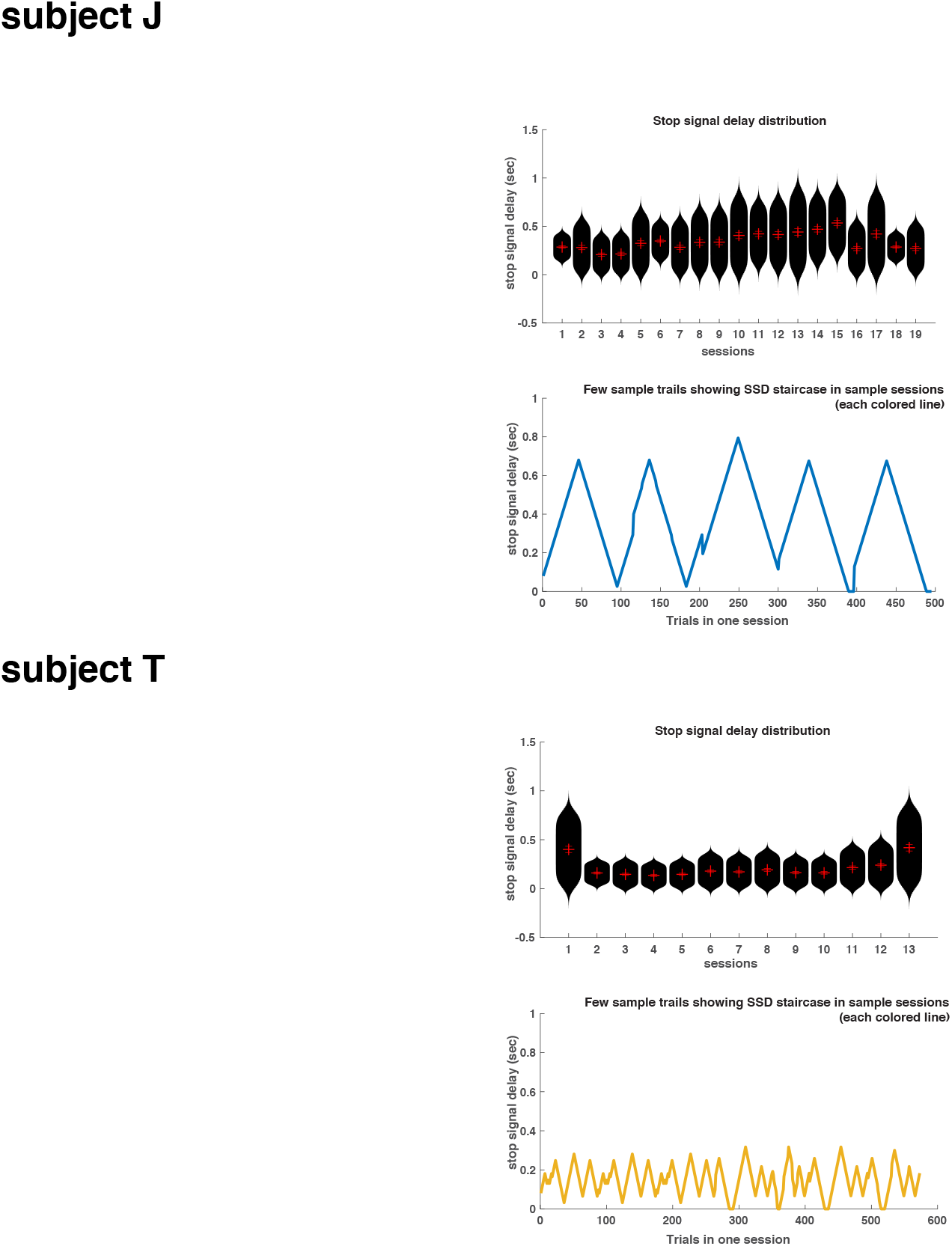

In the next figure, the top panel (A) shows the violin distributions of SSRT for subjects J and T across recorded sessions, whereas the bottom (B) panel draws it out explicitly as a function of session.

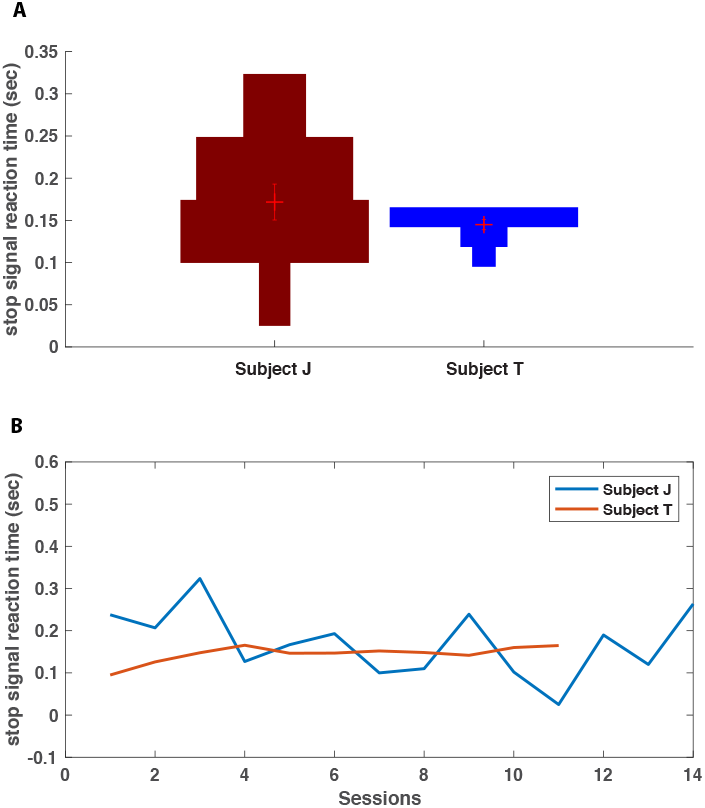

## DATA ACCESSIBILITY

All data will be available on the data section of the Hayden Lab website (www.haydenlab.com/datacode.html, https://doi.org/10.6084/m9.figshare.7865297.v1)

## Acknowledgements

This work was supported by an R01 (DA038615) to BYH. We thank Meghan C. Pesce for help with data collection.

